# Assembly of chromosome-scale and allele-aware autotetraploid genome of the Chinese alfalfa cultivar Zhongmu-4 and identification of SNP loci associated with 27 agronomic traits

**DOI:** 10.1101/2021.02.21.428692

**Authors:** Ruicai Long, Fan Zhang, Zhiwu Zhang, Mingna Li, Lin Chen, Xue Wang, Wenwen Liu, Tiejun Zhang, Long-Xi Yu, Fei He, Xueqian Jiang, Xijiang Yang, Changfu Yang, Zhen Wang, Junmei Kang, Qingchuan Yang

**Affiliations:** Institute of Animal Sciences, Chinese Academy of Agricultural Sciences, Beijing, China; Department of Crop and Soil Sciences, Washington State University, Pullman, WA, USA; School of Grassland Science, Beijing Forestry University, Beijing, China; United States Department of Agriculture-Agricultural Research Service, Plant Germplasm Introduction and Testing Research, Prosser, WA, USA

**Keywords:** Alfalfa, Autotetraploid, Genome assembly, Resequencing, GWAS

## Abstract

Alfalfa (*Medicago sativa* L.), the most valuable perennial legume crop, referred to as “Queen of the Forages” for its high nutritional value and yield production among forage crops. Comprehensive genomic information of germplasm resources from different ecological regions and modern breeding strategies, such as molecular-marker assisted breeding are of great importance to breed new alfalfa varieties with environmental resilience. Here, we report assembly of the genome sequence of Zhongmu-4 (ZM-4), one of the most planted cultivars in China, and identification of SNPs associated with alfalfa agronomic traits by Genome-wide Association Studies (GWAS). Sequence of 32 allelic chromosomes was assembled successfully by single molecule real time sequencing and Hi-C technique with ALLHiC algorithm. About 2.74 Gbp contigs, accounting for 88.39% of the estimated genome, were assembled with 2.56 Gbp contigs anchored to 32 pseudo-chromosomes. In comparison with *M. truncatula* A17, distinctive inversion and translocation on chromosome 1, and between chromosome 4 and 8, respectively, were detected. Moreover, we conducted resequencing of 220 alfalfa accessions collected globally and performed GWAS analysis based on our assembled genome. Population structure analysis demonstrated that alfalfa has a complex genetic relationship among germplasm with different geographic origins. GWAS identified 101 SNPs associated with 27 out of 93 agronomic traits. The updated chromosome-scale and allele-aware genome sequence, coupled with the resequencing data of most global alfalfa germplasm, provides valuable information for alfalfa genetic research, and further analysis of major SNP loci will accelerate unravelling the molecular basis of important agronomic traits and facilitate genetic improvement of alfalfa.

## Introduction

Alfalfa (*Medicago sativa*), one of the most widely planted perennial forage crops in the world, has been considered as the most valuable forage for its high yield production and high nutrient value (Annicchiarico et al., 2015a). Furthermore, alfalfa also has good stress tolerance and nitrogen fixation ability (Li and Brummer, 2012; Redondo et al., 2012). The average alfalfa hay production in America has reached more than 54 million tons per year over the last three years (USDA-crop production, 2020). By the end of 2016, the commercial hay of alfalfa reached 3.79 million tons and the planting area was around 450 thousand ha in China (Shi et al., 2019). As the sharply increasing demand for milk and ruminant animal production in China and other developing countries, the requirement for high quality forage such as alfalfa has becoming a restricted step for animal production supply and quality.

Cultivated alfalfa (*Medicago sativa* ssp. *sativa* L.) is a self-incompatibly cross-pollinated perennial autotetraploid (2n = 4x = 32). The growth and genetic feature of alfalfa makes it difficult to developing an efficient breeding strategy. Recurrent phenotypic selection is still a major and efficient breeding method. However, the method is time consuming and labor-intensive. Marker associate breeding is considered to be an efficient supplemental way for phenotypic selection (Hawkins and Yu, 2018; Li and Brummer, 2012). Along with the developing of marker associate breeding, the genetic architecture of agronomic traits is being reveal. Several candidate SNP markers have been identified for quantitative traits of biomass (Annicchiarico et al., 2015b; Liu and Yu, 2017; Wang et al., 2020), forage quality (Biazzi et al., 2017; Lin et al., 2020; Wang et al., 2016; Yu et al., 2019), and drought/salt tolerance (Yu, 2017; Yu et al., 2016; Zhang et al., 2015). Quantitative trait loci (QTL) associate with yield (Adhikari et al., 2019; McCord et al., 2014; Zhang et al., 2019b) and fall dormancy (Adhikari et al., 2018; Li et al., 2015a) also have been identified. And genomic selection (GS) related study based on alfalfa biomass yield have been conducted (Li et al., 2015b). However, most of these studies were analyzed based on the reference genome of the *M. truncatula*, a close relative of alfalfa (Tang et al., 2014). Beside their high similarity, there are also some distinctive different characters for alfalfa (such as overwintering rate and stress tolerance). Furthermore, some loci were unable to be mapped to *M. truncatula*, suggesting the uniqueness and divergence of alfalfa from its relative specie (Sakiroglu and Brummer, 2017; Yu et al., 2017). A highly-resolved genome is a valuable resource for the marker-based genetic breeding aiming to improve the agronomically important traits of the cultivated alfalfa.

Over a long period in the past, it was a monumental challenge to assemble chromosome-level genome sequences for autopolyploid or highly heterozygous genomes. In 2018, the first allele-aware and chromosome-level genome of autopolyploid *Saccharum spontaneum* was assembled successfully using ALLHiC algorithm developed by Zhang *et al*. (Zhang et al., 2018; Zhang et al., 2019c). It is interesting to note there are two draft genomes of autopolyploid alfalfa, Xinjiangdaye and Zhongmu-1, were reported and provide valuable information for alfalfa related studies recently (Chen et al., 2020; Shen et al., 2020). The genome of Xinjiangdaye alfalfa was assembled to allele-aware chromosome level using ALLHiC algorithm, and the genome of Zhongmu-1 was merely assembled to haploid chromosome level. Compared with other several plants such as maize and rice, the genomic resources of alfalfa are still limited. In this study, we reported an allele-aware and chromosome-level genome sequence assembly of another Chinese alfalfa cultivar Zhongmu-4 (ZM-4), one the most planted alfalfa in Northern China due to its high yield and salt tolerance. In addition, we evaluated 93 agronomic traits of 220 alfalfa germplasm resource accessions from all over the world. Based on our assembled genome sequence and genotyping with whole genome sequencing (WGS), in a total of 111,075 SNP markers were screened. Several SNPs associated with agronomic traits (including salt response and fall dormancy) were identified by genome-wide association study analysis.

## Results

### Genome assembly of autotetraploid alfalfa ZM-4

Flow cytometry assay showed that DNA peak ratio of ZM-4 to Jemalong A17 (*Medicago truncatula*), a close relative of alfalfa with an estimated haploid genome size of 425 Mbp (Tang et al., 2014), was about 3.61 (Supplementary Figure 1), indicating that the tetraploid genome size of ZM-4 is about 3,068 Mbp (Supplementary Table 1), which is close to the estimation of 3,371 Mbp by 2C value (3.44 pg) from Plant DNA C-values database. Using the 168 Gbp Illumina 150 bp pair-end genome sequencing reads, the ZM-4 genome was surveyed and the tetraploid genome size was estimated of 2,962 Mbp by KmerGenie based on *K-mer* frequency (K=19) (Supplementary Figure 2). The estimated genome size of ZM-4 was set as 3.1 Gbp in the following analyses.

A total of 262 Gbp continuous long reads (CLR) were generated from the PacBio Sequel sequencing system (Supplementary Table 2), and used for self-correction and assembly by Canu. About 2.74 Gbp contigs, accounting for 88.39% of the estimated genome size, were assembled with contig N50 of 2.06 Mbp and GC content of 34.2% (Supplementary Table 3). The assembled contig sequences were corrected and polished with 168 Gbp Illumina paired-end sequences, and then preliminarily assembled by 3D-DNA, yielding 49,967 corrected contigs (Supplementary Table 3). The corrected and polished contigs were imported to polyploid genome assembly program ALLHiC to build allele-aware and chromosomal-scale genome using Hi-C paired-end reads. The final assembled genome included 2.56 Gbp contigs anchored to 32 pseudochromosomes (super-scaffolds), which represented 8 groups of homologous chromosomes with 4 sets of monoploid chromosomes, and 0.18 Gbp unanchored contigs. The assembly quality was then assessed by synteny analysis, Hi-C contact matrix, benchmarking universal single-copy orthologs (BUSCO) and the long terminal repeat (LTR) assembly index (LAI). The Hi-C linkage plot indicated that the chromosome groups were clear cut (Figure 1a, Supplementary Table 4). Alignment of the 32 monoploid chromosomes of ZM-4 with *M. truncatula* showed thatthe 4 homologous chromosomes shared high synteny with their counterparts in *M. truncatula*. A translocation was identified between chromosome 4 and 8, and a consistent inversion was observed in chromosome 1 (Figure 1b, c and Supplementary Figure 3). About 98.4% (1,588 out of 1,614) of BUSCOs were detected in the assembled genome (Supplementary Table 5). LAI of whole assembled genome was about 13.85 (among 10~20), indicating our assembly of ZM-4 reaches “Reference” level according to Ou’s classification system (Ou et al., 2018).

**Figure 1.**
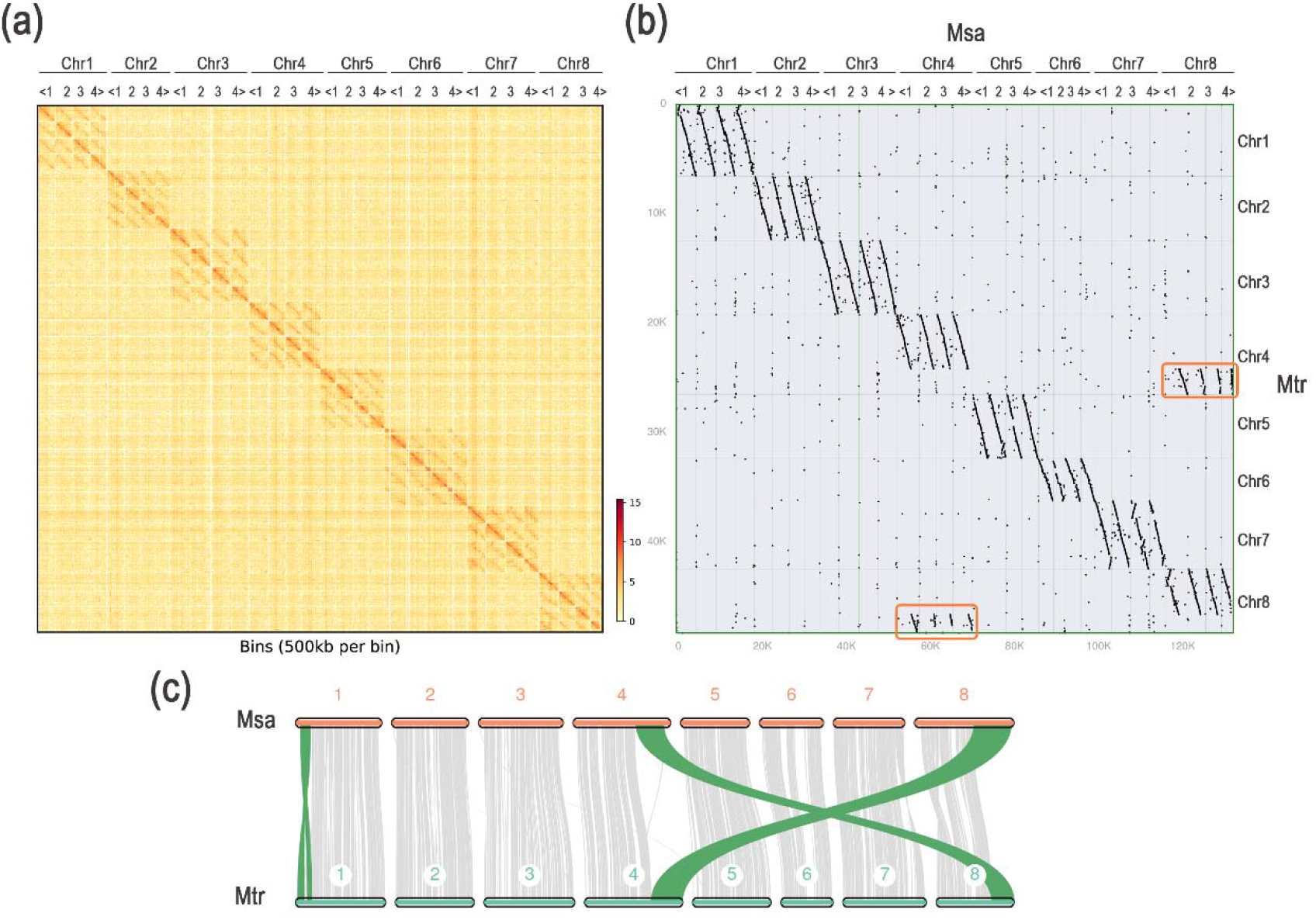
Hi-C linkage plot and synteny dot plot. (a) e Hi-C linkage plot of 32 chromosomes of ZM-4 genome. (b) Alignment of 8 groups of homologous chromosomes with 4 sets of haploid chromosomes of ZM-4 (Msa) with *M. truncatula* (Mtr). (c) Synteny linkage plot of the first set of monoploid chromosomes of ZM-4 (Msa) with *M. truncatula* (Mtr).

A total of 146,704 protein-encoding genes were identified from the assembled allele-aware genome and 93.02% (136,467) genes were functionally annotated by searching in InterPro, EggNOG and KEGG databases. Gene density in the 32 monoploid chromosomes was shown in Figure 2 and Supplementary Table 6.

**Figure 2.**
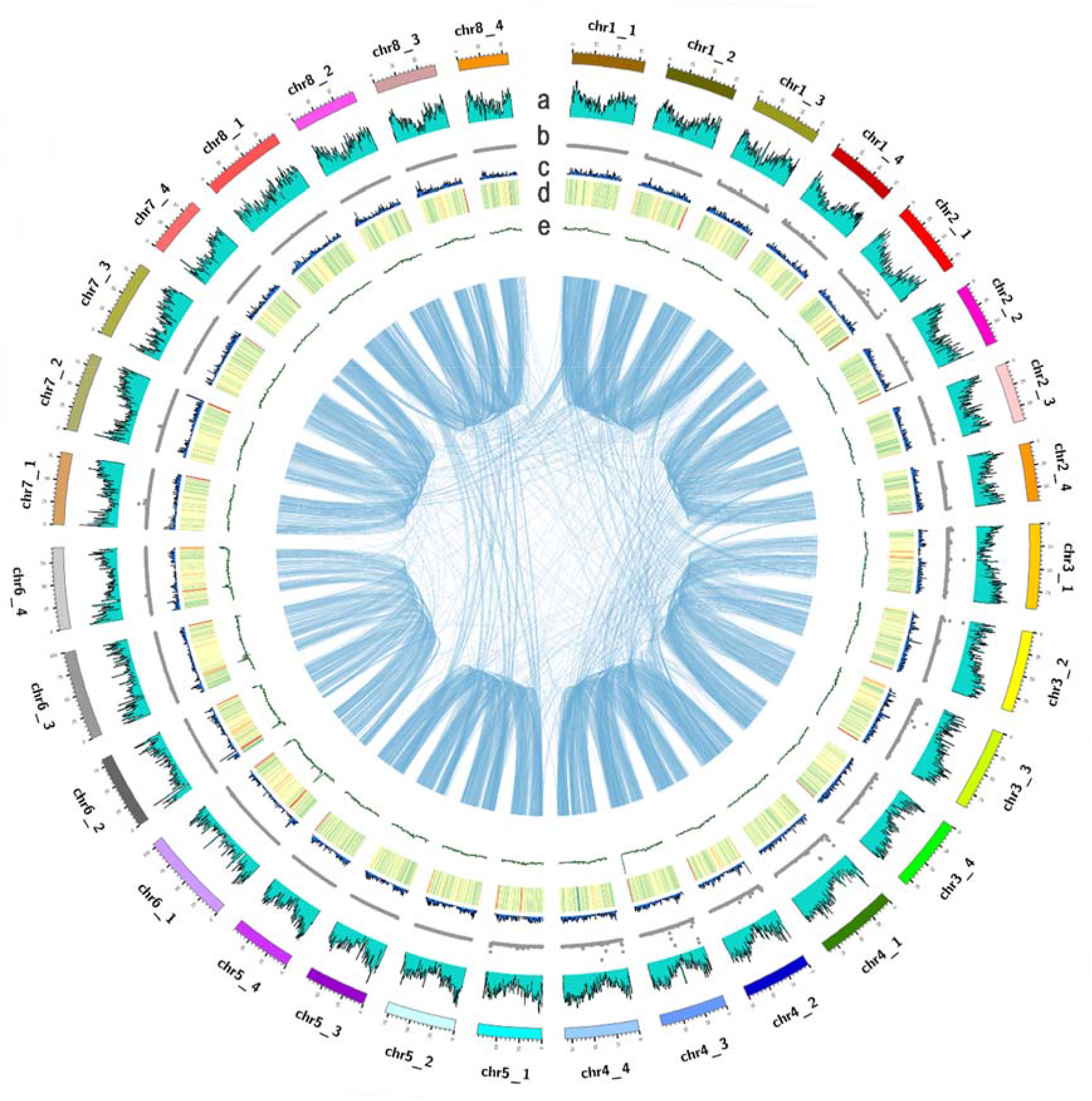
Distribution of genomic features of ZM-4. (a) Density of annotated genes. (b) SNP density among GWAS population. (c) Density of the genes with Ka/Ks ratio > 1 between syntenic paralogous gene pairs. (d) Density of repeat sequences. (e) GC content density. The Links in the center connect the synteny blocks.

### Repeat sequence prediction

A total of 1,558 Mbp repetitive sequences were identified, accounting for 56.80% of the assembled ZM-4 genome (Figure 2), and 49.28% is long terminal repeat (LTR) elements, including Ty1-Copia superfamily retrotransposons (10.93%), Ty3-Gypsy superfamily retrotransposons (18.33%) and unclassified LTR elements (20.02%). The rest repeat sequences are mainly composed of long or short interspersed nuclear elements segments (LINEs or SINEs), DNA elements and simple repeats. In contrast, *M. truncatula* contains 181 Mbp repetitive sequences with 76 Mbp LTR repetitive elements, accounting for 43.94% and 18.36% of the genome, respectively. The proportion of repetitive sequences, especially LTR elements in ZM-4 was significantly larger than that in *M. truncatula*. These results suggested that LTR accumulation is probably one of the main causes for genome enlargement of alfalfa. For individual chromosome, the percentage of repetitive sequences of the sixth chromosomes (Chr6_1~Chr6_4) were much higher than other chromosomes (Figure 2, Supplementary Figure 4). These results, together with the fact that chromosome 6 is the shortest of 8 chromosomes, may explain, to some extent, the relatively lower assembly quality and less protein-encoding genes identified in chromosome 6 (Supplementary Table 6).

### Evolution and whole genome duplication (WGD) analysis

A phylogenetic tree was constructed based on the 28 single-copy orthologous genes identified from *M. sativa* (the first set of ZM-4 chromosomes was used to represent a monoploid alfalfa), *A. thaliana*, *P. trichocarpa* and twelve legume species (Figure 3a). For time calibration, the divergence time of 106 million years (Mya) between *A. thaliana* and *M. sativa* was used as reference. As shown in Figure 3a, *M. sativa* is closer to legume species, especially *M. truncatula*, and the two diverged approximately 5.3 Mya ago. During evolution, gene family size change was observed for the 15 selected species, and a total of 3,377 expanded gene families and 5,626 contracted gene families were found in *M. sativa* (Figure 3a).

**Figure 3.**
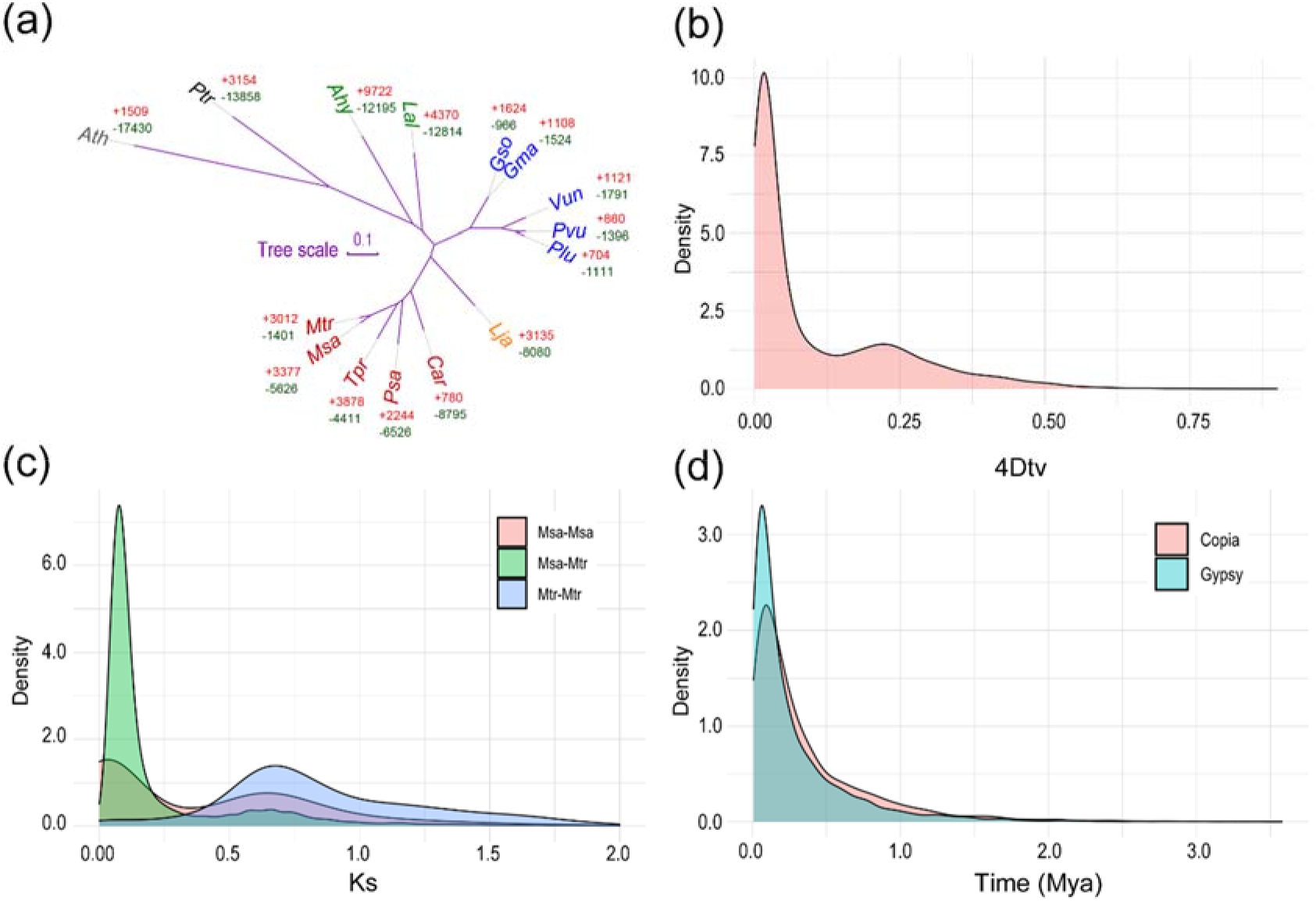
Phylogeny, whole genome duplications and LTR transposable element insertion time distribution. (a) Phylogenetic tree of ZM-4 (Msa) and 14 indicated dicot plant species. “+” and “-” indicate expanded (in red) and contracted gene families (in green), respectively. (b) Density of *4Dtv* value of syntenic paralogous genes in ZM-4. (c) Density of *Ks* value of syntenic paralogous genes in/between ZM-4 (Msa) and *M.truncatula* (Mtr). (d) Density of insert time of Copia and Gypsy LTR transposable elements.

We then analyzed distribution of synonymous substitutions per synonymous site (*Ks*), nonsynonymous substitutions per nonsynonymous site (*Ka*), *Ka/Ks* ratio and four-fold synonymous third-codon transversion (*4Dtv*) of syntenic paralogous genes in ZM-4. Density plot of *4Dtv* and *Ks* value displayed two peaks, which suggested that *M. sativa* has experienced two whole-genome duplication events (Figure 3b, c). Distribution of genes with *Ka/Ks* (between syntenic paralogous gene pairs) > 1 was illustrated in Figure 2c, showing no significant difference between allelic chromosomes. Transposable elements (TEs) play important roles in evolution of genome. The Distribution of LTR transposable elements (TEs) insert time demonstrated that a recent burst of LTR occurred in ZM-4 genome within the last 1.0 Mya (Figure 3d).

### Genotype calling and population structure

To explore SNPs related to agronomic traits of alfalfa, 220 accessions collected from six continents were genotyped by whole genome sequencing (WGS). A total of 29.6 million SNPs were detected using VarScan pipeline. 111,075 SNPs were obtained after data filtration with cutoff of missing value >10%, minimum mean read deep >5 and minor allele frequency (MAF) >0.05. Based on the filtered SNP markers, population structure of subgroup was estimated using admixture with K value from 2 to 10. 220 genotypes displayed a division into three clusters based on CV error (Figure 4a, Supplementary Figure 5). Cluster 1 was dominated by the accessions of Europe. Cluster 2 contained the accessions with diverse background and cluster 3 mainly contained the accessions from Asia (Figure 4b). Principle component analysis (PCA) showed similar results (Figure 4c), and the variation explained by the top three principle components was 3.5%, 1.8% and 1.5%, suggesting a relatively weak population structure of the alfalfa accessions. Linkage disequilibrium (LD) analysis demonstrated that LD decay occurred at 15 kb for the whole genome when *r*^2^ < 0.1 (Figure 4d).

**Figure 4.**
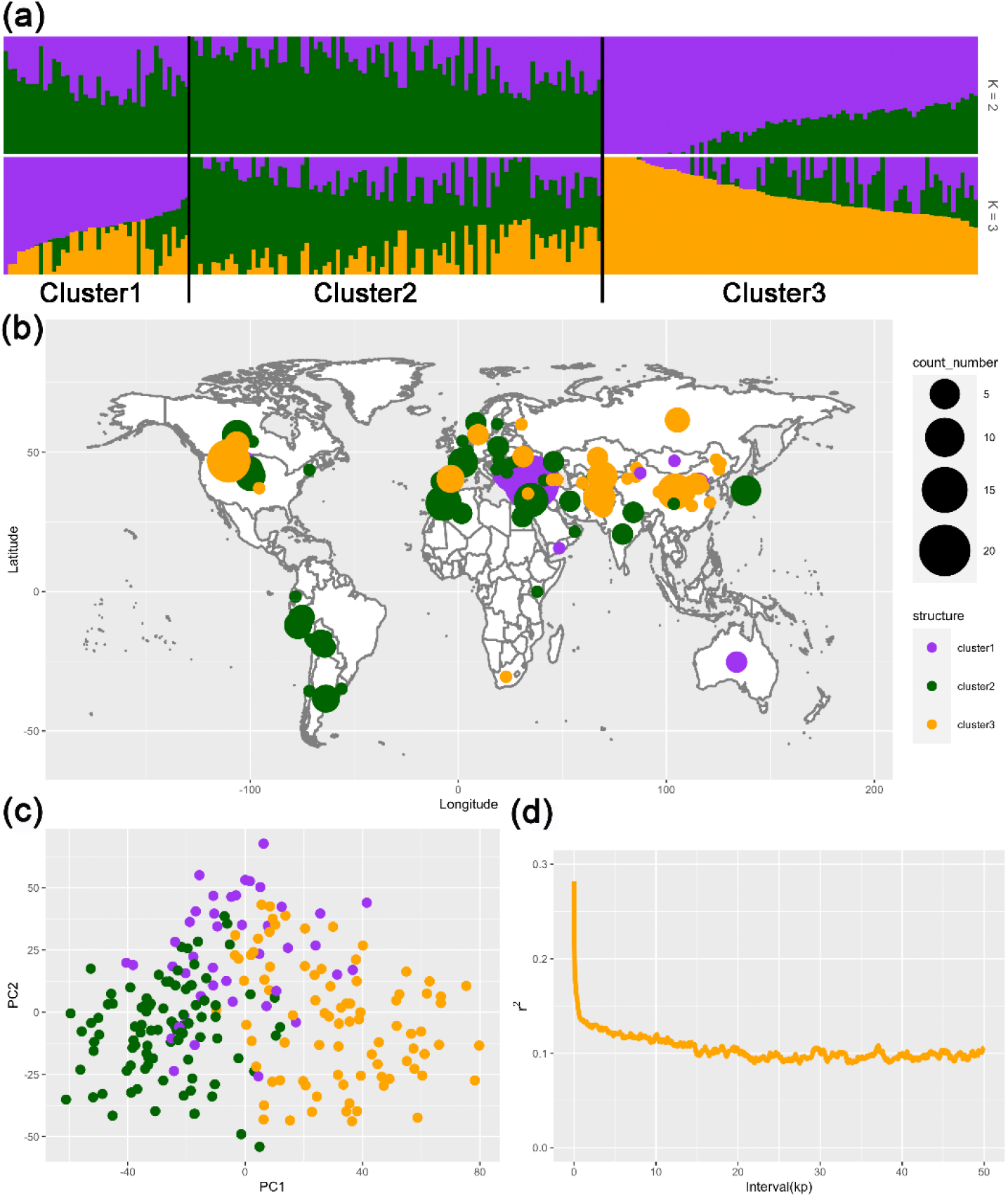
Analysis of population structure and linkage disequilibrium. (a) Population structure shown by cluster plot. (b) Distribution of 220 accessions clustered by population structure on world map. (c) Principal components analysis (PCA) of the PC1 and PC2 pairwise scatter plots. (d) Linkage disequilibrium decay within 50 kb-distance.

### GWAS of phenotypic traits

To identify candidate loci/genes related to important agronomic traits of alfalfa, a total of 93 traits (35 from our lab and 58 from USDA GRIN database) were analyzed by GWAS using the 220 association mapping panel. In total, 101 SNPs were found to be associated with 27 traits, including quality traits (crude protein, neutral detergent insoluble crude protein, et al), disease related traits (aphanomyces, verticillium wilt, et al) and growth-related traits (dormancy height, fall growth, et al). Briefly, the average associated SNP number is 4 for every trait. The trait blue aphid has highest 14 associated SNP. There are 9 traits has one associated SNP (Supplementary Table 7 and Supplementary Figure 6). The trait-associated markers are valuable resource for the marker associated selection in alfalfa breeding.

Several SNPs were detected to be associated with multiple traits. For example, SNP (at 20.41 Mb on Chromosome 3_1) was associated with crude protein (CP), neutral detergent insoluble crude protein (NDICP), Nonfiber Carbohydrates (NFC), and Phosphorus (P), which contribute to alfalfa hay quality/nutrition (Supplementary Table 7, Supplementary Figure 7). We also found an association locus at 67.48 Mb on chromosome 6_2 that was associated with three related traits: dormancy height, fall growth and frost damage (Figure 5a, b, c). Further analysis of allele frequency showed that “A” nucleotide, which was significantly associated with high dormancy, had a higher frequency in the select cultivated alfalfa (0.64) and landrace (0.61) than in wide materials (0.44) (Figure 6a). Geographically, “A” allele had a higher frequency in the cultivars from Asia (0.56), Europe (0.75) and South America (0.8), while the cultivars from North America had a lower frequency (0.25) (Figure 6b). Based on Zhongmu-4 genome, seven protein-encoding genes are annotated within 15 Kb (LD = 15 Kbp, r2 = 0.1) up- and down-stream of the SNP. Expression analysis showed that one candidate gene (Msa0935280) was up-regulated in several genotypes with high dormancy relative to the low dormancy germplasm (Figure 6c). For the SNP (locating at 33.94 Mb on chromosome 4_1) significantly associated with salt stress (Figure 5d), one of the two candidate genes (Msa0552040 and Msa0552050) was up-regulated by 100 mM NaCl and the induction of Msa0552040 was fairly stronger in the salt-tolerant accessions than in the salt-sensitive ones (Figure 6d).

**Figure 5.**
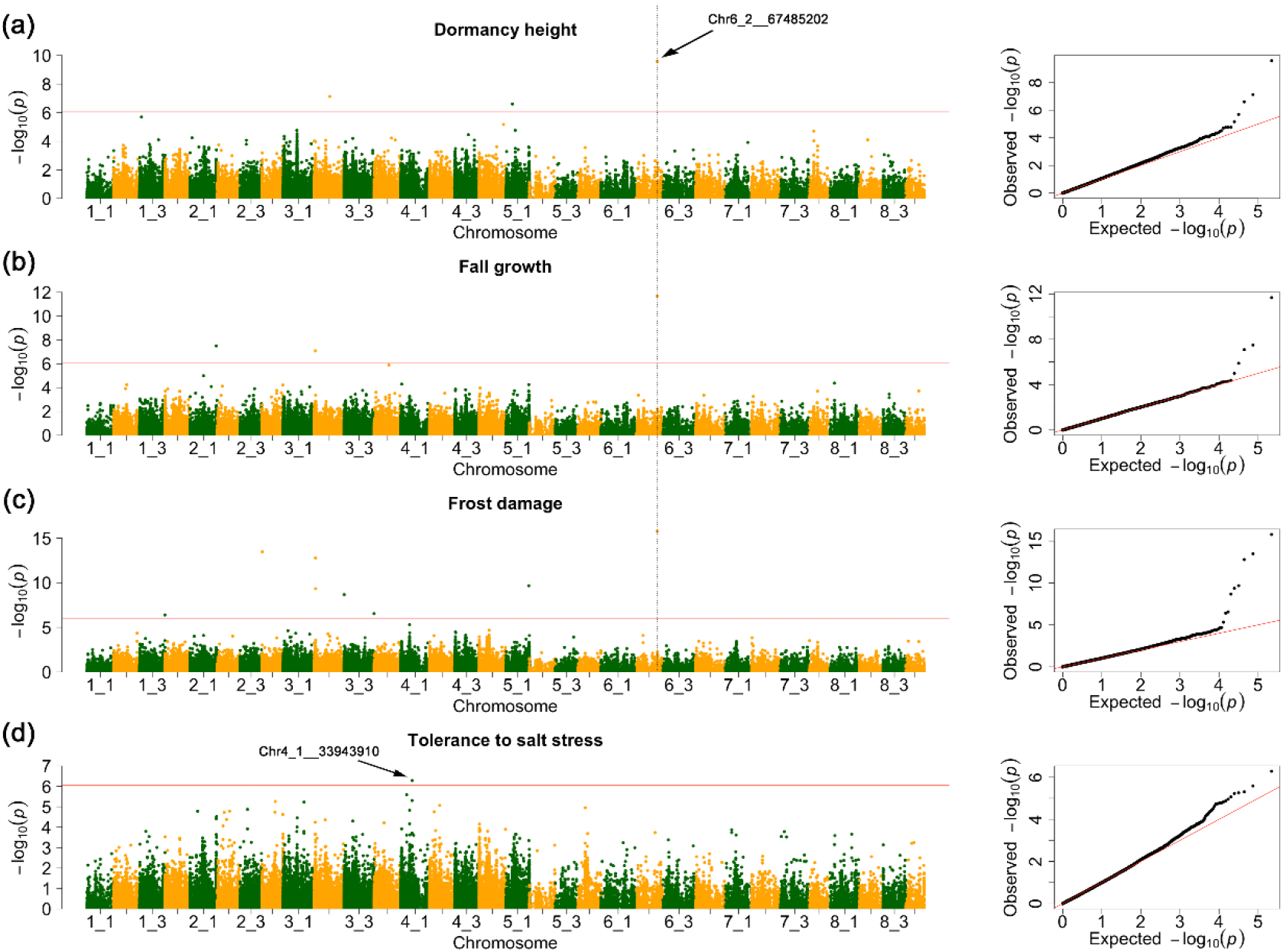
Manhattan plot and Quantile-Quantile (QQ) plot of SNPs associated with fall dormancy and salt tolerance. Manhattan plot (left panel) and QQ plot (right panel) showing SNPs associated with dormancy height (a), fall growth (b) and frost damage (c). Dashed line depicted the location of a SNP simultaneously associated with the three traits and arrow indicated its locus. (d) Manhattan plot (left panel) and QQ plot (right panel) showing SNPs associated with trait of salt-responsiveness. GWAS was performed by GAPIT3 software with the function of Blink and significant p-value threshold set at P= 9.00×10^-7^(0.1/111075).

**Figure 6.**
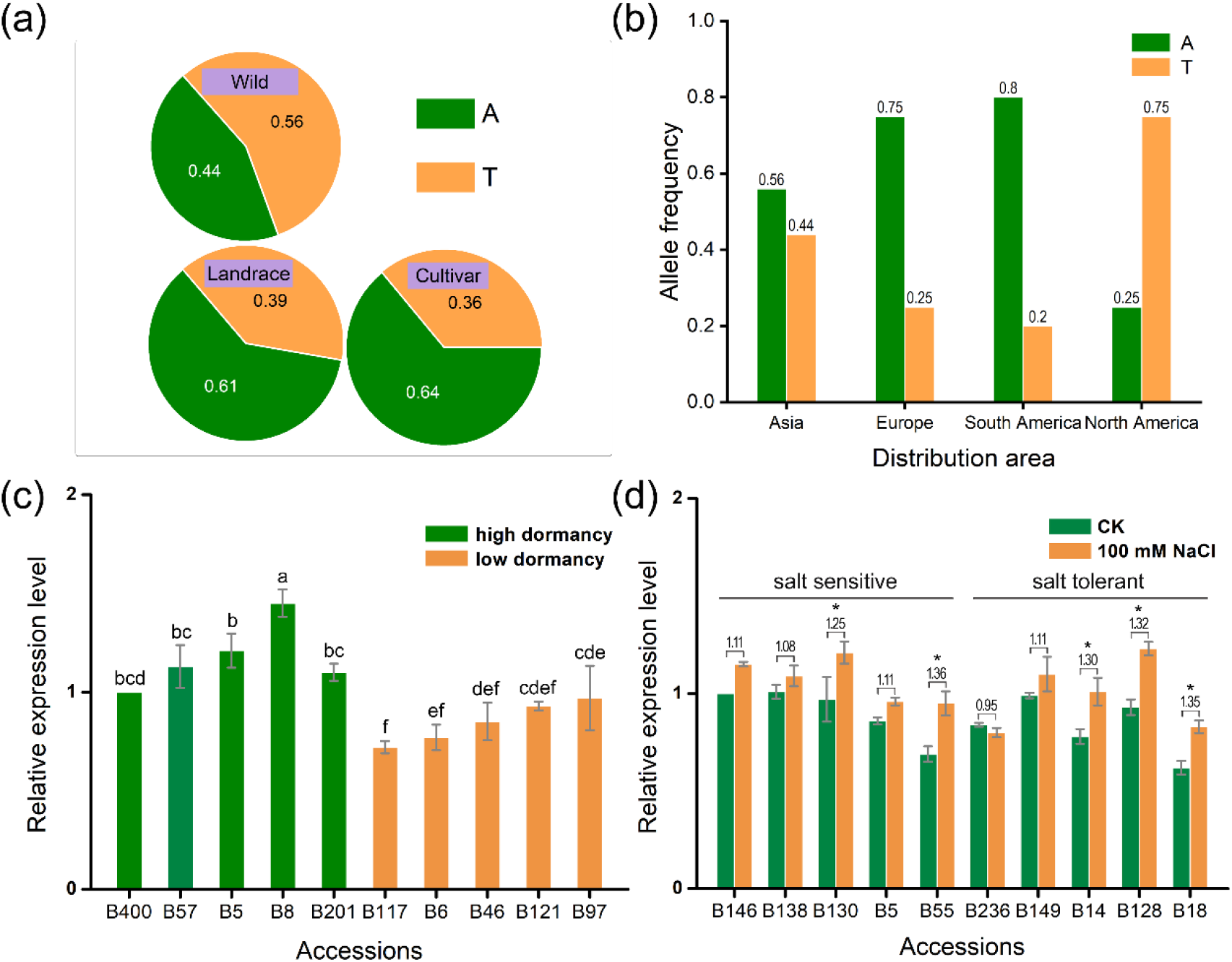
Analysis of allele frequency of a multi-effect SNP and the relative expression of two candidate genes in the representative accessions. (a) Allele frequency of SNP (Chr6_2_67485202) in wide materials, landrace and cultivated alfalfa. (b) Allele frequency of Chr6_2_67485202 in cultivars from Asia, Europe, South and North America. (c) Analysis of the relative expression level of a candidate gene for SNP (Chr6_2_67485202) in the representative accessions, and data were normalized arbitrarily to one accession (on left). (d) The relative expression of *BRI1 SUPPRESSOR 1 (BSU1)-LIKE 3* in the representative accessions sensitive or tolerant to salt stress.

## Discussion

### Assembly of autotetraploid alfalfa genome

This study reported a reference genome of a Chinese autotetraploid alfalfa cultivar Zhongmu-4 by exploiting the third-generation sequencing, Hi-C sequencing and allele-aware chromosome level assembly algorithm. Besides Xinjiangdaye alfalfa, this is the second allele-aware and chromosome-level reference genome of autotetraploid alfalfa (Chen et al., 2020). Recently Shen et al. also reported an assembly of chromosome-level genome sequence of our alfalfa cultivar Zhongmu-1, the genome of which was only assembled to monoploid-level (Shen et al., 2020). Because of the outcrossing and self-incompatibility feature, the genome and genotype may be distinctly different between different alfalfa cultivars and individuals. Accordingly, we assembled an allele-aware chromosome level reference genome of Zhongmu-4, a representative alfalfa cultivar of Zhongmu series. Compared with other several plants such as maize and rice, the genome resources of alfalfa are still limited. Our study provides a new genome to the growing list of sequenced alfalfa genomes. For a long time, the studies of molecular biology, evolution, GWAS and QTL for alfalfa were restricted for the lack of reference genome. Traditional alfalfa improvement breeding requires long time and screening of large population. Molecular marker-assisted breeding approach based on high-quality reference genome can shorten breeding cycle and increase the breeding efficiency. To examine the genome quality and identify agronomic traits associating molecular markers, we used the assembled genome of Zhongmu-4 as the reference genome for population structure and GWAS analysis of 220 accessions.

### The genetic diversity of alfalfa

The improvement of alfalfa is majorly based on recurrent phenotypic selection and outcrossing. The genetic exchange among different alfalfa individuals always exists in alfalfa growing regions (Li and Brummer, 2012). It is more difficult to distinguish alfalfa accessions based on geography origin than self-pollination plants, such as soybean and rice (Huang et al., 2012; Zhou et al., 2015). Our results showed that the top three PCs only explained 6.8% genetic variance. That means weak population structure was exist among alfalfa. However, based on the population structure cluster information, the accessions comping from different geography have some part of genetic segregation. For example, the accessions coming from Asia and South America can be separate into different genetic clusters (Figure 4b). This phenomenon was also found in another alfalfa core germplasm population (Shen et al., 2020). One possible reason is that alfalfa accessions may experience different domestication and improvement process to adapt different geographic environment. But the outcrossing nature and breeding strategy generating strong gene flow between different accessions. The adapted genes for different region can be exchanged and help alfalfa spread widely in the world. Furthermore, the accessions comping from Europe have widely genetic diversity. This result may be the reason that alfalfa origin from the Caucasus region: northeastern Turkey, Turkmenistan and northwestern Iran. This phenomenon can also be detected in previous studies (Qiang et al., 2015; Sakiroglu et al., 2010; Wang et al., 2020).

### The GWAS results of alfalfa

Because the population structure is weak for alfalfa. It is relatively easier to control false positive SNP compared to self-pollination crops, such as rice (Huang et al., 2010) and soybean (Fang et al., 2017). It can be found in our QQ plot result (Figure5 and Supplemental figure6). This feature makes alfalfa no need to construct artificial population to control population structure. However, heterozygote is a bigger problem to do GWAS for alfalfa. We detected SNP markers base on single allele chromosome, our results still showed that 68% markers have more the 10% heterozygous genotype among individuals. These heterozygous genotype makes it harder to estimate an accurate minor allele frequency (MAF) and filter MAF markers. Furthermore, the suppose of GWAS method is that most SNP is homozygous genotype among individuals. If there are too many heterozygote, the GWAS results may be disturb by these markers (Jia et al., 2017; Wang et al., 2020; Yu et al., 2017). Although there are some new featured to do GWAS for alfalfa, we still identified 101 significant associated SNP among 27 phenotypes. The further analysis of one SNP (chr6_2_67485202) showed that different allele frequency was found among cultivar/landrace and wild accessions. And further analysis of cultivar showed that different geography has different allele frequency. These results showed a similar domestication and improvement pattern with other crops, such as maize (Guo et al., 2018) and soybeans (Zhou et al., 2015). Further analysis the candidate gene (Msa0935280) of SNP chr6_2_67485202 showed a different expression level among high and low dormancy accessions. This gene may play a key role for alfalfa dormancy. Another one salt stress gene showed a smaller difference among different salt tolerance groups. But its homologous gene has been proved to be associated with salt tolerance in *Arabidopsis* (Cui et al., 2013). The function of this gene in alfalfa need to be further validated. Taken together, the SNP identified by us can reveal some genetic information of alfalfa and provide some useful SNP marker for molecular breeding and candidate gene studies.

## Methods

### Plant materials and genome sequencing

A Chinese native alfalfa cultivar Zhongmu-4 (*Medicago sativa* L. cv. Zhongmu-4), one of the most planted alfalfa in North China for its high yield, nutritive value, fast regeneration, and resilience to environmental stress, was sequenced for genome assembly. Fresh leaves, stems and flowers were collected from one five-year-old individual plant of Zhongmu-4 alfalfa (ZM-4/Msa) in Langfang planting base (39.59°N, 116.59°E). DNA was extracted from leaves using PlantZol Kit (TransGen Biotech, China). A portion of DNA was constructed 20-kb continuous long reads (CLR) libraries and sequenced using PacBio Sequal platform (Pacific Biosciences, USA). In total, 19 Single Molecule Real Time (SMRT) cells were sequenced. The other portion DNA was constructed pair-end libraries and sequenced by Illumina NovaSeq platform (Illumina, USA). The sequencing was done by Biomarker Technologies Corporation (China) and Novogene Corporation (China).

A total of 220 alfalfa accessions, including 55 cultivar, 26 cultivated material, 95 landrace, 4 breeding material, 16 wild material and 23 uncertain improvement status materials (Supplementary Table 8), were used in genome-wide association study.

Among them, 26 accessions are obtained from the database of Medium Term Library of National Grass Seed Resources of China and 194 accessions are obtained from the database of U.S. National Plant Germplasm System. The accessions were mainly collected from China, the United States, Turkey, Afghanistan, Tashkent Uzbekistan, Spain, Russia, Morocco, France and Argentina. In 2018, the individuals of 220 accessions were planted on the field of the Chinese Academy of Agricultural Sciences (CAAS) in Langfang, Hebei Province, China. A randomized complete block design with three replications was used for the experimental design. Every accession included 5 individuals and separated with each other with 30 cm in a single row for one replication. The accessions spacing was 65 cm between rows and column. In order to keep all individuals were uniform, they were clipped to a height of 5 cm after transplant. Furthermore, all individuals were clipped again at the early flowering stage (when 10% of plants begin flowering). A total of 93 phenotype traits were used in the GWAS analysis, including 4 agronomic traits (fall growth, dormancy, leaf long and leaf wide), 31 nutrition traits and 58 phenotype traits from USDA GRIN database (31 growth related phenotype and 27 resistance related phenotype, https://npgsweb.ars-grin.gov/gringlobal/AccessionObservation.aspx?id=1340810) (Supplementary Table 9).

### Hi-C library construction and sequencing

Hi-C libraries were constructed as described in previous report (Wang et al., 2015). Fresh and young leaves of ZM-4 were fixed with formaldehyde. The denatured chromatin was isolated from the fixed leaves and digested with restriction enzyme HindIII. The restriction ends were labeled with biotin and ligated to form chimeric junctions. The chimeric fragments were isolated and then constructed to five Hi-C libraries that were sequenced on an Illumina NovaSeq platform (Illumina, USA) with 2×150 bp pair-end reads. A total of 951 million read pairs (285 Gb) were produced.

### Genome estimation and assembly

The genome size of ZM-4 was estimated by *K-mer* frequency-based method with KmerGenie (v1.7051) using Illumina pair-end sequencing reads (Chikhi and Medvedev, 2014). The genome size also measured by flow cytometry. Fresh leaves from ZM-4 and *M. truncatula* L. cv. Jemalong A17 (a close homology plant with complete sequencing genome of 425 Mbp) were chopped finely with a sharp blade in 250 μl Cystain PI Absolute P (Sysmex, Japan) nuclei extraction buffer. After incubation for 60 sec, the homogenate was filtered through 50 μm strainer. The nuclei filtrate was mixed with 500 μl staining buffer (containing 50 μg·ml^-1^ propidium iodide and 5 U·ml^-1^ RNase A) and incubated for 15 min at room temperature under dark condition. The fluorescence intensities of nuclear DNA were measured on LSRFortessa platform (BD Biosciences, USA). The genome size of ZM-4 was assessed by comparing the PI means with reference Jemalong A17. The genome size also was estimated according to Plant DNA C-values database (https://cvalues.science.kew.org/).

The genome was assembled as following strategy: (1) PacBio CLR subreads were assembled to contigs using Canu (v1.8) with default parameters (Koren et al., 2017). The genome size was set as 3000 Mbp. (2) The obtained contigs were polished by genome Illumina data using Pilon (v1.23) and then interrupted by Juicer (v1.5) according to the Hi-C data and then preliminarily assembled to scaffolds by 3D-DNA (v180419) (Dudchenko et al., 2017; Durand et al., 2016). (3) The obtained scaffolds were assembled to chromosomes with a chromosomal-scale autopolyploid genome assembly program ALLHiC (Zhang et al., 2018; Zhang et al., 2019c). The detailed procedures of this step were followed by the instruction of ALLHiC on github (https://github.com/tangerzhang/ALLHiC). The gene annotation data of *Medicago truncatula Mt4.0v1* from Phytozome database (v12) was used for chromosome assembly and collinearity analysis (Tang et al., 2014). GMAP (v2013-10-28) was used to generate an allelic contig table to remove noisy Hi-C signals (Wu and Watanabe, 2005). MCScanX was employed to draw gene synteny plot (Wang et al., 2012).

### Gene annotation

RNA-Seq data and protein sequences of homology species *Medicago truncatula* (Mt4.0v1) were used for ab initio gene prediction. The RNA samples extracted from the leaves, stems and flowers were reverse transcribed to cDNA, constructed pair-end libraries and then sequenced by Illumina NovaSeq platform. RNA-Seq data were assembled by Trinity (v2.9.0) with default parameters of de novo and genome-guided assembly pipelines (Grabherr et al., 2011). Two rounds of gene annotation running were carried out. The assembled transcripts and homology protein sequences were firstly annotated by BRAKER (v2.15), which allowed for fully automated training of the gene prediction tools GeneMark-EX (v4.61) and AUGUSTUS (v3.3.3) from RNA-Seq and/or homology protein sequence (Bruna et al., 2020; Hoff et al., 2019; Stanke et al., 2008). In the second round of annotation, the previously trained GeneMark and AUGUSTUS files, assembled transcripts and homology proteins were imported into MAKER (v2.31.10) pipeline (http://www.yandell-lab.org/software/maker.html) to integrate transcript evidence and protein-coding evidence. BUSCO (v4) was used for evaluation of annotation completeness (Simao et al., 2015).

### Repeat sequence prediction

A de novo transposable element (TE) sequence library of the assembled genome was firstly built by RepeatModeler (v2.0.1, http://www.repeatmasker.org/RepeatModeler/), which can automatically employ three de novo repeat-sequence annotation programs (RECON, v1.08, RepeatScout, v1.0.6 and LTR_retriever (v2.8.7)) to identify repeat elements from genome sequence. The repeat sequence library customized by RepeatModeler were then imported to RepeatMasker (v4.0.9, http://www.repeatmasker.org/) to identify and cluster repetitive elements. Tandem Repeat Finder (v 4.09) was used to find tandem repeats (Benson, 1999). To further identify LTRs, LTR_FINDER (v1.07) and LTRharvest (a module of genometools v1.5.10) were employed to investigate LTRs respectively (Ellinghaus et al., 2008; Gremme et al., 2013; Xu and Wang, 2007). LTR_retriever (2.8.7) was use to integrate results of LTR_FINDER and LTRharvest. LTR assembly index (LAI) was also calculated by LTR_retriever, which was used to assess the assembly quality of genome (Ou et al., 2018).

### Phylogeny and gene family evolution

The protein sequences of first set of chromosomes (Chr x_1) of ZM-4 and 14 other plant species (*Lotus japonicus*, *Pisum sativum*, *Arabidopsis thaliana*, *Arachis hypogaea*, *Cicer arietinum*, *Glycine max*, *Glycine soja*, *Lupinus albus*, *Medicago truncatula*, *Phaseolus lunatus*, *Phaseolus vulgaris*, *Trifolium pratense*, *Vigna unguiculata*, *Populus trichocarpa*) were used for phylogenetic tree analysis by OrthoFinder (v2.3.11) (Emms and Kelly, 2015). Based on the specie phylogenetic tree and the divergence time (http://timetree.org/) of *Arabidopsis thaliana* and *Medicago sativa* (~106 Mya), a ultrametric tree (timetree) was obtained by r8s program (v1.81). Then the changes in gene family size in these species were analyzed by CAFE (v4.2.1) (Han et al., 2013).

### Synteny and whole genome duplication (WGD) analysis

The syntenic blocks between each monoploid of ZM-4, as well as between monoploid of ZM-4 with *M. truncatula* were identified by MCScanX (Wang et al., 2012). *Ka* (the ratio of the number of nonsynonymous substitutions per nonsynonymous site) and *Ks* (the ratio of the number of synonymous substitutions per synonymous site) values of syntenic paralogous genes were calculated by PERL script “add_ka_and_ks_to_collinearity.pl” of MCScanX. The 4Dtv (transversion of four-fold degenerate site) values of syntenic paralogous genes were calculated by PERL script “calculate_4DTV_correction.pl” (https://github.com/JinfengChen/Scripts/blob/master/FFgenome/03.evolution/distance_kaks_4dtv/bin/calculate_4DTV_correction.pl).

### Gene function annotation and classification

The protein-coding genes of ZM-4 were functionally annotated by eggNOG-mapper (v 5.0) (Huerta-Cepas et al., 2019) and InterProScan (v5.36-75.0) (Zdobnov and Apweiler, 2001). Kyoto Encyclopedia of Genes and Genomes (KEGG) pathways of annotated genes were analyzed by KEGG automatic annotation server of KAAS (Moriya et al., 2007).

### Resequencing and SNP calling

The young leaves (two week after regrowth) of one representative individual was chosen from each accession for resequencing. Total DNA was extracted using the CWBIO Plant Genomic DNA Kit (CoWin Biosciences, Beijing, China), according to the manufacturer’s protocol. At least 6 ug of genomic DNA from each accession was used to construct a sequencing library following the manufacturer’s instructions (Illumina Inc.). Paired-end sequencing libraries were sequenced by BerryGenomics company on Illumina NovaSeq6000 platform (Illumina, USA) with 2*150 bp pair-end reads. About 10 Gb Sequencing data with average Q30 of 85% were produced for each accession. Paired-end sequencing reads were mapped to the assembled genome of ZM-4 with BWA-MEM using default parameters (Li, 2013). Samtools was used to translate SAM file to BAM file and sort BAM files using default parameters (Li et al., 2009). Picard was used mark duplicate reads and GenomeAnalysisTK were used to correct indels which can be mistaken as SNP. Samtools mpileup and VarScan were used to detect SNP (Koboldt et al., 2012). Furthermore, SNP data were filtered using Vcftools (Danecek et al., 2011) with missing rate less than 10%, minor allele frequency more than 0.05 and mean reads deep more than 5.

### Population structure and linkage disequilibrium (LD)

Population structure was calculated using the software of admixture with default parameter (Alexander et al., 2009) and the best K value was generated by the value of CV error. Principal components analysis (PCA) was calculated using GAPIT3 (Wang and Zhang, 2018) with the function of principal components analyses. The PCA results combined with geography information were showed using R package ggplot2 (Ginestet, 2011). LD information was calculated using the software of PopLDdecay with default parameters (Zhang et al., 2019a). The VCF file containing all 111,075 SNP markers information were import to PopLDdecay. The LD results among all accessions were used to estimate LD of alfalfa. The distance against mean R^2^ within 50 kb of LD were showed using R package ggplot2.

### Association mapping

GWAS were conducted using GAPIT3 with the function of Blink (Huang et al., 2019; Wang and Zhang, 2018). The first three PCA values were used as fixed effects in the model to correct false positive. The significant SNP threshold was bonferroni multiple test threshold (P=0.1/111,075= 1.78×10^-7^). The Blink method use iterations to select a set of markers associated with a trait. These associated markers are fitted as covariate for testing the rest markers. This method has a better control on false positives than kinship approach. The real data and simulated data results showed that Blink has a higher statistical power than other method, such as MLM, FarmCPU et al (Huang et al., 2019). The Manhattan plots and QQ plots of GWAS results were showed using the R package qqman (Turner, 2014).

### qRT-PCR verification of associated genes

Associated genes were selected based on the up and down stream candidate region of LD decay distance of significant SNP. Alfalfa accessions with the top five maximum or the minimum values evaluated by the standard of tolerance to salt or fall growth and frost damage were selected for the gene expression test. Seeds were imbibed at 4°C for two days then moved to long-day conditions with 16 h light (24°C)/8 h dark (20°C). The seedlings were kept for 4 days before transferring to Hoagland solution (half strength) to grow 10 days. For salt treatment, 14-day-old seedlings were supplemented with NaCl (150 mM), and the shoots were collected after growing for 3.5 days (84 h) under LD. The leaves of 5 accessions with high dormancy ratings and 5 accessions with low dormancy ratings were sampled in October for fall dormancy associated gene expression analysis. Total RNA was extracted using the Mini BEST Plant RNA Extraction Kit (Takara Biotech Co., Ltd., Dalian, China) and cDNA was synthesized using the one-step PrimeScript RT-PCR Kit (Takara) according to the manufacturer’s instructions. Quantitative PCR was performed in an ABI7300 system (Applied

Biosystems, Foster City, CA, USA) using SYBR Premix Ex Tap II kit (Takara). Data represent the calculation based on the mean threshold cycle (Ct) values of three biological replicates. Alfalfa *β-actin* gene was used as reference.

## Data availability

The data supporting the results of this article are included within the article and the provided Supplementary files. The genome assembly and gene annotation files available at https://figshare.com/s/fb4ba8e0b871007a9e6c. The genome and transcriptome sequencing raw data described in this article are publicly available in the NCBI database under project PRJNA685277. The population resequencing data was uploaded to the National Genomics Data Center (NGDC, https://bigd.big.ac.cn/) under BioProject PRJCA004024. The genome annotation data of *Lotus japonicus* was downloaded from http://www.kazusa.or.jp/lotus/. The genome annotation data of *Pisum sativum* was downloaded from https://urgi.versailles.inra.fr/Species/Pisum. The genome annotation data of *Arabidopsis thaliana, Arachis hypogaea, Cicer arietinum, Glycine max, Glycine soja*, *Lupinus albus*, *Medicago truncatula*, *Phaseolus lunatus*, *Phaseolus vulgaris*, *Trifolium pratense*, *Vigna unguiculata*, *Populus trichocarpa* were downloaded from Phytozome (https://phytozome-next.jgi.doe.gov/).

## Acknowledgements

This work was supported by the National Natural Science Foundation of China (grants 31971758 to Q. Yang and 32071865 to R. Long), Collaborative Research Key Project between China and EU (2017YFE0111000), the earmarked fund for China Agriculture Research System (CARS-34), the Agricultural Science and Technology Innovation Program (ASTIP-IAS14).

## Author contributions

R. Long, Q. Yang and J. Kang conceived and designed the research. R. Long performed genome assembly, genome annotation and evolution analysis. F. Zhang and L. Chen performed the GWAS analysis. R. Long, F. Zhang, T. Zhang, F. He, X. Jiang, X. Yang, C. Yang prepared samples and performed phenotyping. X. Wang, W. Liu and Z. Wang performed the gene expression analysis. R. Long, F. Zhang, Z. Wang, J. Kang, M. Li, L. Chen, L. Yu, Z. Zhang, and Q. Yang wrote and revised the paper. All authors read and approved the final manuscript.

## Supporting information

Additional supporting information may be found online in the Supporting Information section at the end of the article.

**Supplementary Figure 1** The PI-A value of *M. truncatula* Jemalong A17 and ZM-4 measured by flow cytometry.

**Supplementary Figure 2** K-mer frequency analysis result (K=19).

**Supplementary Figure 3** The synteny linkage plot of set2, set3, set4 chromosomes of ZM-4 with chromosomes of *M. truncatula* and the synteny linkage plot within set1, set2, set3, set4 chromosomes.

**Supplementary Figure 4** The repetitive sequences percentage of 32 chromosomes and whole genome of ZM-4.

**Supplementary Figure 5** The cross-validation (CV) error plot from k=1 to k=10. The best value of k=3 clusters which is the value with lowest CV error.

**Supplementary Figure 6** The quantile-quantile (QQ) plot of 27 agronomic traits with significant associated SNP. The trait name is on the top of QQ plot.

**Supplementary Figure 7** Manhattan plot of seven protein related traits. X-axis showed chromosome name.

**Supplementary Table 1** The PI-A value of ZM-4 alfalfa (Msa) and *Medicago truncatula* (Mtr) measured by flow cytometry and the estimated genome size of ZM-4.

**Supplementary Table 2** The sequencing data information (the whole tetraploid genome size of ZM-4 was set as 3.1 Gbp).

**Supplementary Table 3** The Canu assembled contigs and Hi-C corrected contigs.

**Supplementary Table 4** Summary of the assembled genome of ZM-4.

**Supplementary Table 5** The benchmarking in BUSCO notation for the assembled genome and annotated proteins of ZM-4.

**Supplementary Table 6** Gene annotation result summary.

**Supplementary Table 7** The associated SNP information of 27 traits.

**Supplementary Table 8** The basic information of 220 alfalfa accessions.

**Supplementary Table 9** The phenotypic information of 93 agronomic traits.

