## Supplementary figure and table for "Assembly of chromosome-scale and allele-aware autotetraploid genome of the Chinese alfalfa cultivar Zhongmu-4 and identification of SNP loci associated with 27 agronomic traits": Supplementary figure 1-7.docx

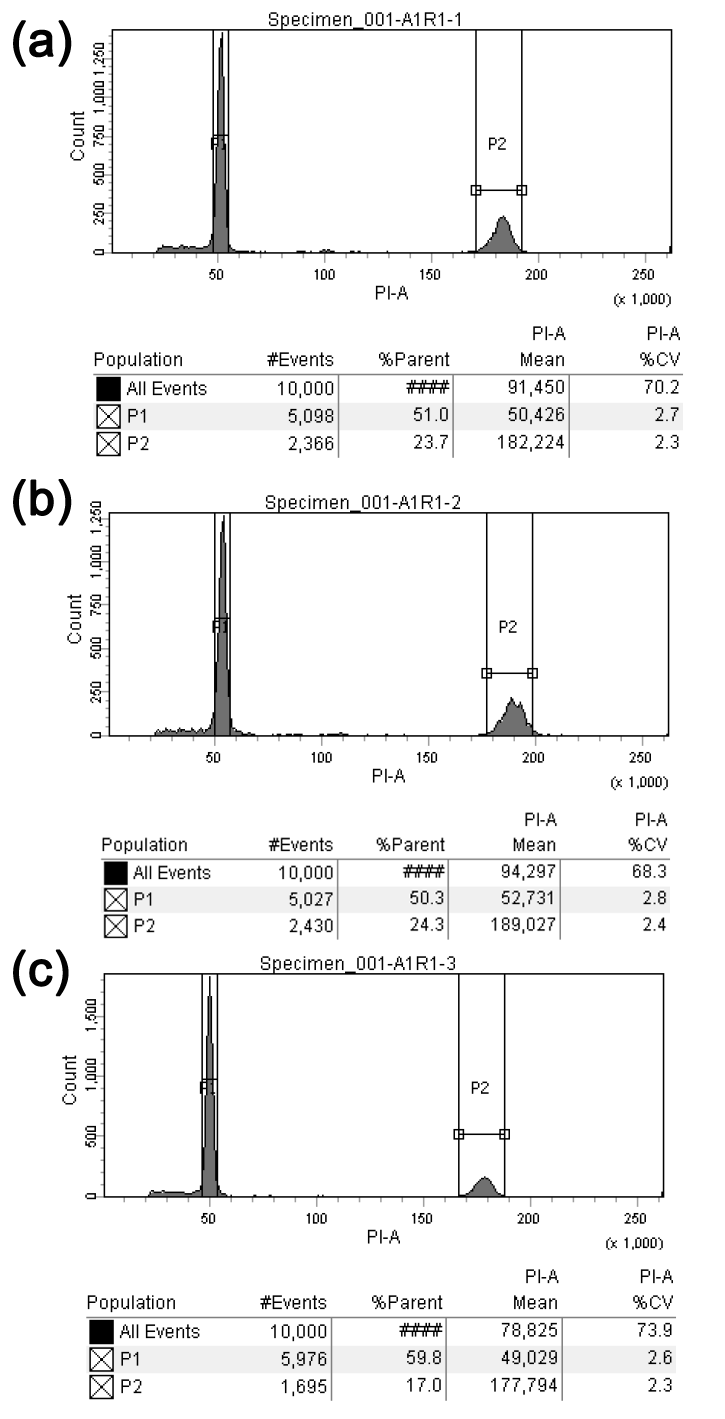

**Supplementary Figure 1.** The PI-A value of *M. truncatula* Jemalong A17 and ZM-4 measured by flow cytometry. (a), (b), (c) represent the results of three independent repeats. P1 and P2 represent Jemalong A17 and ZM-4 respectively. The average of PI-A ratio (ZM-4/ Jemalong A17) is about 3.61.

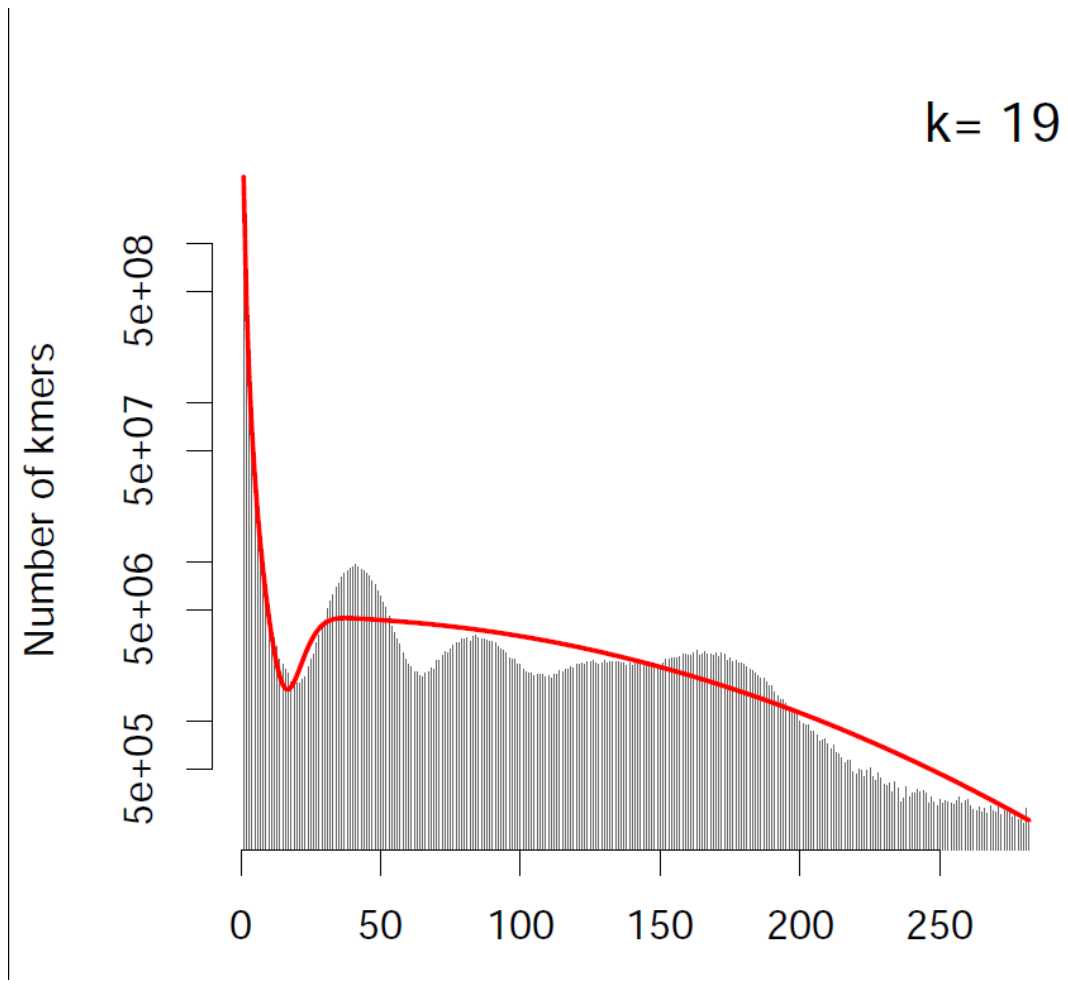

**Supplementary Figure 2.** *K*-mer frequency analysis result (K=19). The haploid genome size of ZM-4 was estimated as 1,480,963,752 bp by KmerGenie, and thus the estimated whole genome size was about 2962 Mbp (1,480,963,752 bp × 2).

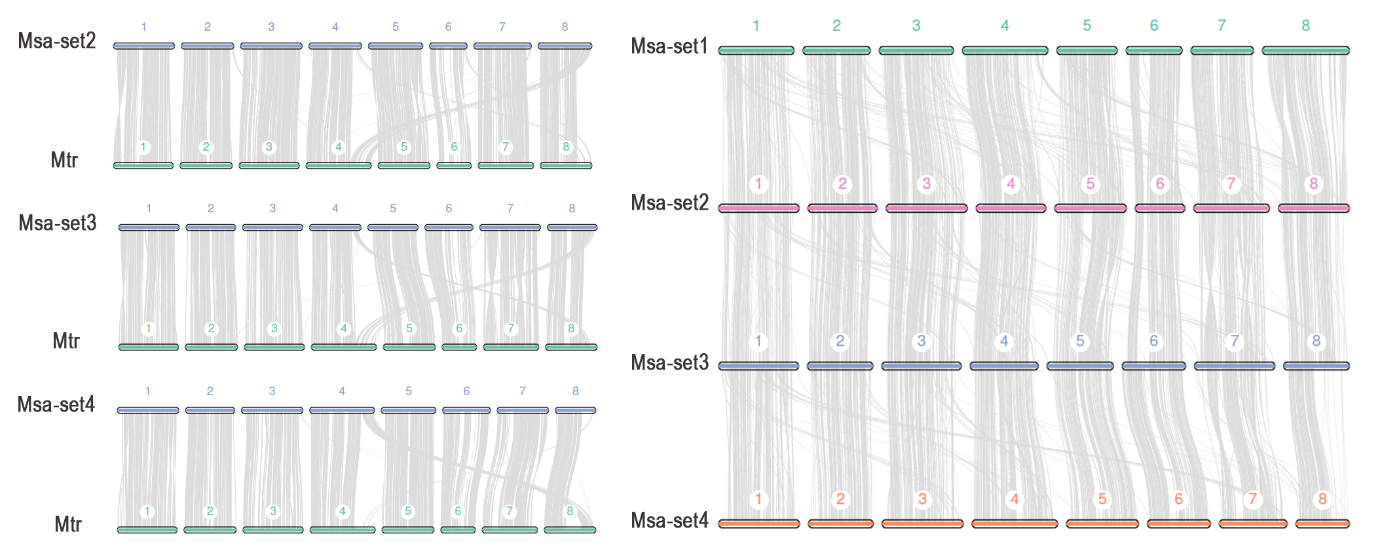

**Supplementary Figure 3.** The synteny linkage plot of set2, set3, set4 chromosomes of ZM-4 with chromosomes of *M. truncatula* and the synteny linkage plot within set1, set2, set3, set4 chromosomes.

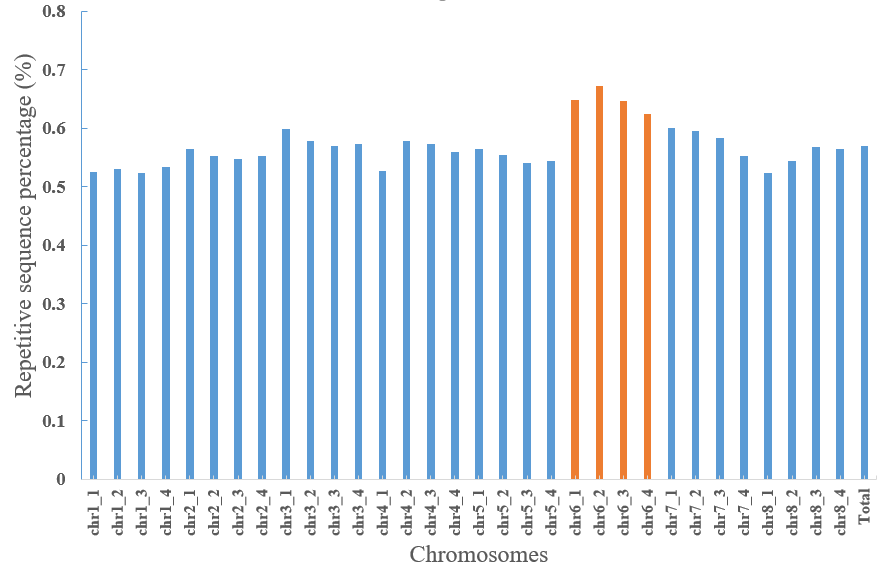

**Supplementary Figure 4**. The repetitive sequences percentage of 32 chromosomes and whole genome of ZM-4.

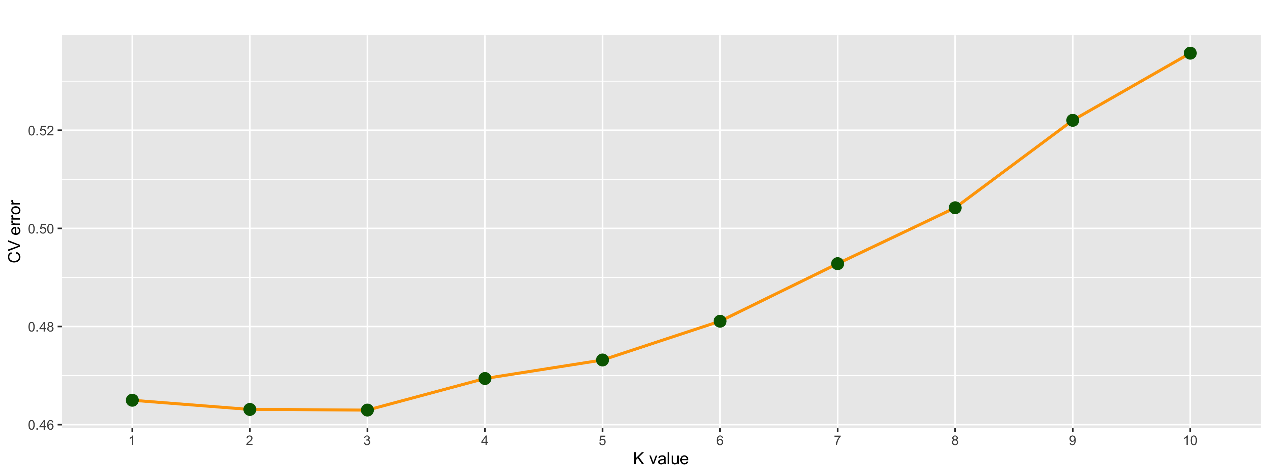

**Supplementary Figure 5.** The cross-validation (CV) error plot from k=1 to k=10. The best value of k=3 clusters which is the value with lowest CV error.

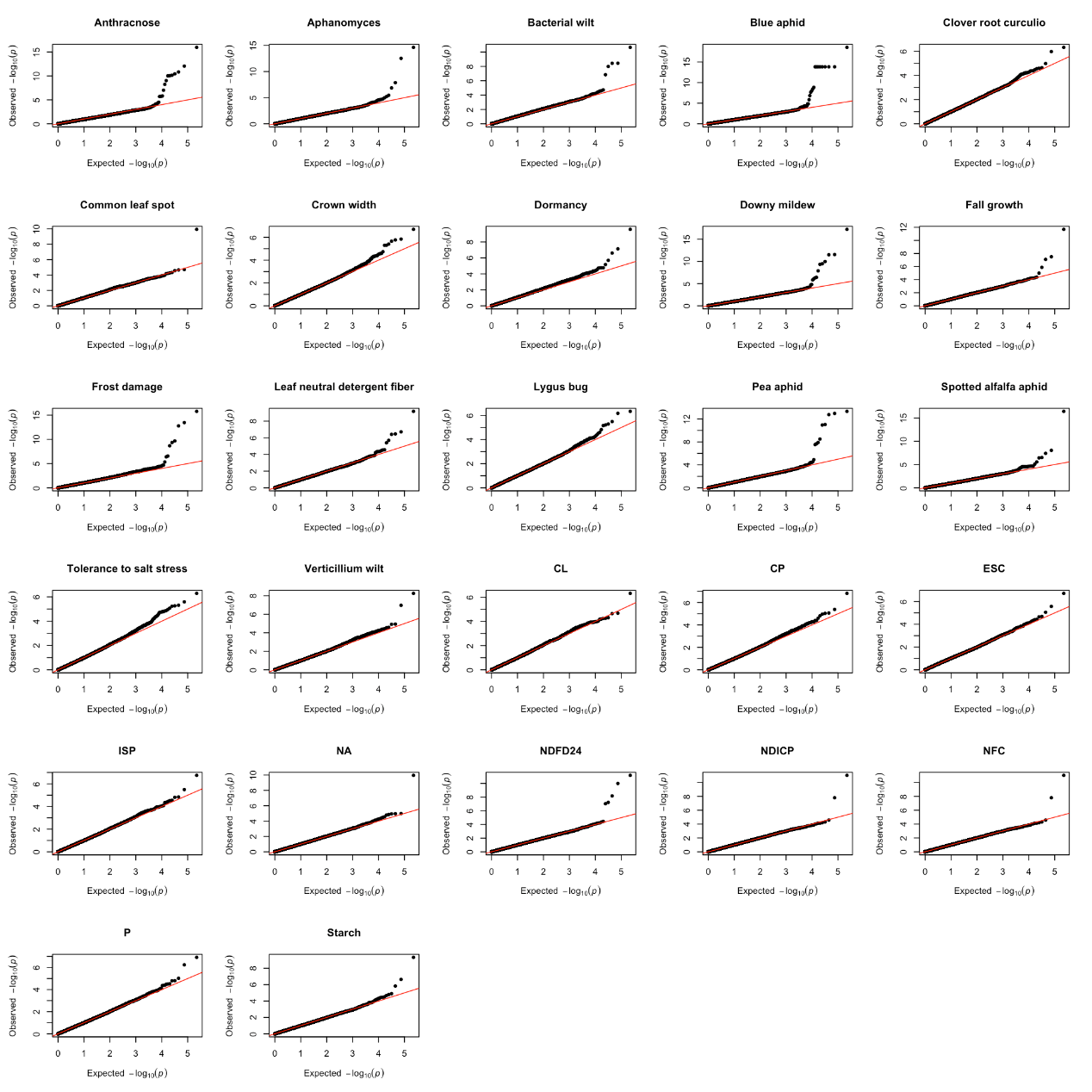

**Supplementary Figure 6**. The quantile-quantile (QQ) plot of 27 agronomic traits with significant associated SNP. The trait name is on the top of QQ plot.

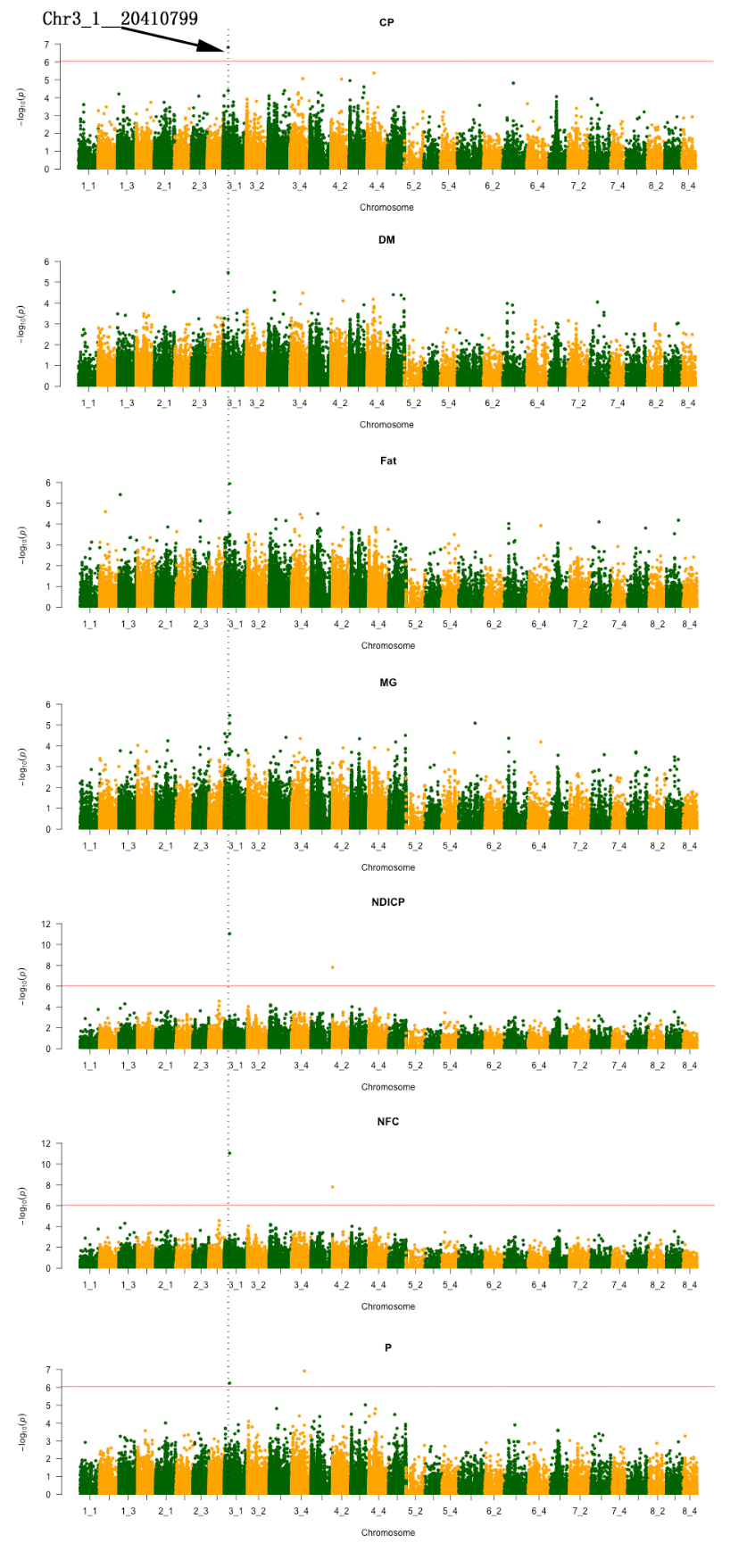

**Supplementary Figure 7**. Manhattan plot of seven protein related traits. X-axis showed chromosome name. They are assigned from 1_1 to 8_4. Y-axis represent the -log_10_(p) value. The black dish line represents the significant SNP. CP, Crude Protein. DM, Digestible Energy. Fat, Fat of alfalfa. MG, Magnesium. NDICP, Neutral Detergent Insoluble Crude Protein. NFC, Nonfiber Carbohydrates. P, Phosphorus.
