## Supplementary figure and table for "Assembly of chromosome-scale and allele-aware autotetraploid genome of the Chinese alfalfa cultivar Zhongmu-4 and identification of SNP loci associated with 27 agronomic traits": Supplementary Table 1-6.docx

**Supplementary Table 1. The PI-A value of ZM-4 alfalfa and *Medicago truncatula* (Mtr) measured by flow cytometry and the estimated genome size of ZM-4**

| **Repeat** | **PI-A(ZM-4)** | **PI-A(Mtr)** | **Foldchange of PI-A ratio** | **Average foldchange of PI-A ratio** | **Monoploid genome size of Mtr (Mbp)** | **Estimated** **tetraploid** **genome size of ZM-4 (Mbp)** |
| --- | --- | --- | --- | --- | --- | --- |
| **Repeat 1** | 177794 | 49029 | 3.63 | 3.61 | 390~425 | 2814~3068 |
| **Repeat 2** | 189027 | 52731 | 3.58 |  |  |  |
| **Repeat 3** | 182224 | 50426 | 3.61 |  |  |  |

**Supplementary Table 2. The sequencing data information (the whole tetraploid genome size of ZM-4 was set as 3 Gbp).**

| **Data type** | **Data size (Gbp)** | **Depth (×)** | **N_50_ (bp)** |
| --- | --- | --- | --- |
| PacBio CLR | 262 | 85 | 29,708 |
| Illumina read (genome) | 168 | 54 | 150 |
| Illumina read (Hi-C) | 285 | 92 | 150 |
| Illumina read (transcriptome) | 26 | - | 150 |

**Supplementary Table 3. The Canu assembled contigs and Hi-C corrected contigs.**

| **Contig** | **Number of contig** | **Total length (Gbp)** | **N50 (kbp)** | **GC (%)** |
| --- | --- | --- | --- | --- |
| Canu assembled contigs | 5000 | 2.74 | 2060 | 34.20 |
| Hi-C corrected contigs | 49967 | 2.74 | 94 | 34.20 |

**Supplementary Table 4. Summary of the assembled genome of ZM-4.**

| **Category** | **Contig number** | **Length (bp)** |
| --- | --- | --- |
| chr1_1 | 1405 | 81475151 |
| chr1_2 | 1466 | 79614812 |
| chr1_3 | 1358 | 77152361 |
| chr1_4 | 1439 | 76635544 |
| chr2_1 | 1335 | 82498830 |
| chr2_2 | 1141 | 70773673 |
| chr2_3 | 1139 | 66739748 |
| chr2_4 | 1190 | 65143524 |
| chr3_1 | 1678 | 94727156 |
| chr3_2 | 1662 | 91622233 |
| chr3_3 | 1796 | 91676550 |
| chr3_4 | 1483 | 82148590 |
| chr4_1 | 1537 | 87806598 |
| chr4_2 | 1277 | 76139727 |
| chr4_3 | 1321 | 74558508 |
| chr4_4 | 1455 | 82903785 |
| chr5_1 | 1052 | 71963113 |
| chr5_2 | 1529 | 78992507 |
| chr5_3 | 1166 | 69949269 |
| chr5_4 | 1275 | 70080956 |
| chr6_1 | 1938 | 104280083 |
| chr6_2 | 1538 | 83446217 |
| chr6_3 | 1875 | 99942479 |
| chr6_4 | 1789 | 91272922 |
| chr7_1 | 1174 | 81116394 |
| chr7_2 | 1528 | 88039813 |
| chr7_3 | 1577 | 87612856 |
| chr7_4 | 1110 | 64180167 |
| chr8_1 | 1311 | 91953106 |
| chr8_2 | 1186 | 73305697 |
| chr8_3 | 1252 | 69667704 |
| chr8_4 | 1267 | 56218238 |
| unanchored contigs | 4864 | 179289811 |
| Total contigs | 49967 | 2742928122 |
| Anchor rate (%) | 93.45 | |

**Supplementary Table 5. The benchmarking in BUSCO notation for the assembled genome and annotated proteins of ZM-4.**

| **Search mode** | **Complete BUSCOs** | **Fragmented BUSCOs** | **Missing BUSCOs** | **Total searched BUSCOs** |
| --- | --- | --- | --- | --- |
| assembled genome | 1588(98.4%) | 5(0.3%) | 21(1.3%) | 1614 |
| annotated proteins | 1552(96.1%) | 17(1.1%) | 45(2.8%) | 1614 |

**Supplementary** **Table 6. Gene annotation result summary.**

| **category** | **total length** | **annotated gene number** | **cds length** | **cds length percent** | **cDNA length** | **cDNA length percent** | **gene length** | **gene length percent** |
| --- | --- | --- | --- | --- | --- | --- | --- | --- |
| chr1_1 | 81475151 | 4808 | 5693277 | 6.99% | 7129491 | 8.75% | 19561034 | 24.01% |
| chr1_2 | 79614812 | 4729 | 5737290 | 7.21% | 7166829 | 9.00% | 19579851 | 24.59% |
| chr1_3 | 77152361 | 4626 | 5409660 | 7.01% | 6745112 | 8.74% | 18527290 | 24.01% |
| chr1_4 | 76635544 | 4415 | 5305050 | 6.92% | 6621100 | 8.64% | 18348694 | 23.94% |
| chr2_1 | 82498830 | 4288 | 4963065 | 6.02% | 6138066 | 7.44% | 17188537 | 20.83% |
| chr2_2 | 70773673 | 4060 | 4685250 | 6.62% | 5761616 | 8.14% | 16059709 | 22.69% |
| chr2_3 | 66739748 | 3787 | 4434615 | 6.64% | 5500904 | 8.24% | 15482882 | 23.20% |
| chr2_4 | 65143524 | 3531 | 4128648 | 6.34% | 5167416 | 7.93% | 14746191 | 22.64% |
| chr3_1 | 94727156 | 4727 | 5614509 | 5.93% | 6959131 | 7.35% | 19189192 | 20.26% |
| chr3_2 | 91622233 | 4766 | 5684358 | 6.20% | 7058504 | 7.70% | 19411778 | 21.19% |
| chr3_3 | 91676550 | 4598 | 5524530 | 6.03% | 6833366 | 7.45% | 18922820 | 20.64% |
| chr3_4 | 82148590 | 4427 | 5283189 | 6.43% | 6502009 | 7.91% | 17798884 | 21.67% |
| chr4_1 | 87806598 | 5446 | 6500250 | 7.40% | 8263854 | 9.41% | 22625002 | 25.77% |
| chr4_2 | 76139727 | 4089 | 4796601 | 6.30% | 6021828 | 7.91% | 16238061 | 21.33% |
| chr4_3 | 74558508 | 3938 | 4639266 | 6.22% | 5839379 | 7.83% | 16303257 | 21.87% |
| chr4_4 | 82903785 | 4668 | 5571273 | 6.72% | 6993784 | 8.44% | 18989225 | 22.91% |
| chr5_1 | 71963113 | 3847 | 4238202 | 5.89% | 5304540 | 7.37% | 14792582 | 20.56% |
| chr5_2 | 78992507 | 4212 | 4750587 | 6.01% | 5917249 | 7.49% | 16482853 | 20.87% |
| chr5_3 | 69949269 | 3836 | 4403655 | 6.30% | 5494350 | 7.85% | 15393684 | 22.01% |
| chr5_4 | 70080956 | 3930 | 4416939 | 6.30% | 5545328 | 7.91% | 15349274 | 21.90% |
| chr6_1 | 104280083 | 3563 | 3805755 | 3.65% | 4581904 | 4.39% | 13354937 | 12.81% |
| chr6_2 | 83446217 | 2906 | 3050577 | 3.66% | 3662576 | 4.39% | 10773400 | 12.91% |
| chr6_3 | 99942479 | 3669 | 3903099 | 3.91% | 4706576 | 4.71% | 14558121 | 14.57% |
| chr6_4 | 91272922 | 3437 | 3602808 | 3.95% | 4331539 | 4.75% | 13033427 | 14.28% |
| chr7_1 | 81116394 | 3978 | 4636509 | 5.72% | 5735609 | 7.07% | 15987591 | 19.71% |
| chr7_2 | 88039813 | 4448 | 5106495 | 5.80% | 6301766 | 7.16% | 17297382 | 19.65% |
| chr7_3 | 87612856 | 4649 | 5330322 | 6.08% | 6655169 | 7.60% | 18612750 | 21.24% |
| chr7_4 | 64180167 | 3698 | 4350816 | 6.78% | 5380309 | 8.38% | 14669171 | 22.86% |
| chr8_1 | 91953106 | 5540 | 6691755 | 7.28% | 8403948 | 9.14% | 23365355 | 25.41% |
| chr8_2 | 73305697 | 4167 | 4973262 | 6.78% | 6212684 | 8.48% | 17503687 | 23.88% |
| chr8_3 | 69667704 | 3802 | 4475643 | 6.42% | 5529124 | 7.94% | 15744350 | 22.60% |
| chr8_4 | 56218238 | 2911 | 3592509 | 6.39% | 4464161 | 7.94% | 12604053 | 22.42% |
| unanchored contigs | 179289811 | 13208 | 14623239 | 8.16% | 18102100 | 10.10% | 45185928 | 25.20% |
| total | 2742928122 | 133496 | 155299764 | 5.66% | 192929221 | 7.03% | 538495024 | 19.63% |
